# Mass-spectrometry-based near-complete draft of the *Saccharomyces cerevisiae proteome*

**DOI:** 10.1101/2020.06.24.168526

**Authors:** Yuan Gao, Lingyan Ping, Duc Duong, Chengpu Zhang, Eric B. Dammer, Yanchang Li, Peiru Chen, Lei Chang, Huiying Gao, Junzhu Wu, Ping Xu

## Abstract

Proteomics approaches designed to catalogue all open reading frames (ORFs) under a defined set of growth conditions of an organism have flourished in recent years. However, no proteome has been sequenced completely so far. Here we generate the largest yeast proteome dataset, including 5610 identified proteins using a strategy based on optimized sample preparation and high-resolution mass spectrometry. Among the 5610 identified proteins, 94.1% are core proteins, which achieves near complete coverage of the yeast ORFs. Comprehensive analysis of missing proteins in our dataset indicate that the MS-based proteome coverage has reached the ceiling. A review of protein abundance shows that our proteome encompasses a uniquely broad dynamic range. Additionally, these values highly correlate with mRNA abundance, implying a high level of accuracy, sensitivity and precision. We present examples of how the data could be used, including re-annotating gene localization, providing expression evidence of pseudogenes. Our near complete yeast proteome dataset will be a useful and important resource for further systematic studies.

## Introduction

Mass spectrometry (MS) is widely applied for protein identification in recent decades. Development of the related technologies, including improved sample preparation, mass spectrometers, as well as downstream bioinformatics analysis, have helped to improve protein identification accuracy and coverage (Domon & Aebersold, 2006; Kumar & Mann, 2009; Mallick & Kuster, 2010; Shevchenko *et al*, 1996b; Tyanova *et al*, 2016; Washburn *et al*, 2001). MS-based proteomics is a powerful tool to obtain high quality measures of the proteome, greatly contributing to our understanding about the composition and dynamics of subcellular organelles, protein interaction, protein posttranslational modification as well as signaling networks regulation (Choudhary & Mann, 2010; Domon & Aebersold, 2006; Jensen, 2006; Pandey & Mann, 2000). However, due to various analytical limitations (Gstaiger & Aebersold, 2009; Nilsson *et al*, 2010; Vanderschuren *et al*, 2013), achieving high quantification accuracy and complete proteome coverage remains a challenge.

*Saccharomyces cerevisiae*, one of the most extensively characterized model organisms, has been subjected to the most comprehensive proteome-wide investigations, including global and organelle-specific proteome (de Godoy *et al*, 2008; de Godoy *et al*, 2006; Ghaemmaghami *et al*, 2003; Ho *et al*, 2018; Huh *et al*, 2003; Kolkman *et al*, 2006; Nagaraj *et al*, 2012; Picotti *et al*, 2009; Picotti *et al*, 2013; Reinders *et al*, 2006; Wiederhold *et al*, 2009; Zahedi *et al*, 2006). The first large-scale proteomic study on yeast has identified 150 proteins (Shevchenko *et al*, 1996a). Later, the number of identified proteins increased to thousands. Specifically, two studies expressing tandem affinity purification(TAP) tag (Ghaemmaghami *et al*., 2003) or GFP tag (Huh *et al*., 2003) in yeast gene natural chromosomal location show that as much as 4500 proteins are expressed during normal growth condition. Subsequent emerging targeted proteomics workflows (Deutsch *et al*, 2008; King *et al*, 2006; Kuster *et al*, 2005), by gathering as many as available yeast MS-based proteomics datasets to construct high quality and coverage protein lists, have substantially improved the yeast proteome to a higher coverage. Complementary absolute quantitative proteomics experiments further validate the expression levels (de Godoy *et al*., 2008; Nagaraj *et al*., 2012). Ho et al. (2018) combined 21 quantitative yeast proteome datasets, including MS-, GFP- and western blotting-based methods, to generate an unified protein abundance dataset, covering about 5400 proteins (Ho *et al*., 2018). This number is still lower than the number of currently annotated 6717 yeast ORFs in SGD database. Moreover, the protein abundance identified solely based on MS is known to span multiple orders of magnitudes, ranging from 2^5^ to 2^21^ copies per yeast cell (Picotti *et al*., 2009). This suggests that many low-abundance proteins have not yet been detected (de Godoy *et al*., 2006). Based on a high-throughput peptide synthesis technique, Picotti et al. (2013) generated an almost completed theoretical yeast proteome, covering 97% of the genome-predicted proteins (Picotti *et al*., 2013). However, the synthesized peptides were artificially selected for favorable MS properties and uniqueness and do not accurately reflect endogenous peptides that would be generated by experimental conditions on actual samples. So this large dataset represents a theoretical result, and may be more valuable for the development and optimization of computational methods.

Despite the challenges, recent technical and methodological developments keep emerging, enabling the almost complete quantitative *Arabidopsis* proteome (Mergner *et al*, 2020) and human proteome draft (Kim *et al*, 2014; Wilhelm *et al*, 2014), which provide useful resources for further function analysis. It also encourages us to look into the possibility of complete coverage of yeast proteome. In this study, we combine the optimized sample preparation (extensive gel molecular weight fractionation, and two digestion enzymes) and a more sensitive and faster liquid chromatography/tandem mass spectroscopy (LC-MS/MS) platform (Orbitrap Velos coupled to a nanoAcquity UPLC), providing the largest yeast proteome dataset to date. In total, we identify 5610 proteins, covering 83.5% annotated yeast ORFs. Among, our dataset shows nearly complete coverage of core proteins, up to 94.1%. We find that proteins are missed mainly due to physical properties, such as small protein molecular weight, high sequence similarity, as well as absence in transcription and uncharacterized gene function. Quantitative analysis of our proteome shows that protein abundance spans six orders of magnitudes, and highly correlate with mRNA abundance, suggesting the high coverage and sensitive of our dataset. Moreover, systematic analysis shows our proteome covers 98% of the annotated KEGG pathways, providing insight into the expression pattern of yeast at the molecular level. Also, we use a select sample to show how this near complete yeast proteome can be used to reannotate the yeast genome.

## Results

### Generation of a deep-coverage yeast proteome with high reliable protein identification

To develop methods for the high coverage proteomics analysis, we started with in-gel digestion coupled with mass spectrometric analysis strategy (GeLC-MS/MS) for the separation and identification of the yeast total cell lysate (TCL) samples cultured in the yeast extract peptone dextrose (YPD) medium (Fig 1A). Firstly, SDS-PAGE was used to resolve the samples, resulting in clear and sharp bands, which indicated the proteins were extracted and separated in high quality and resolution (Fig 1B). Each lane was excised into 26 gel bands based on the molecular weight (MW) and the protein abundance. The proteins in these gel bands were in-gel digested with trypsin or endoproteinase LysC (lysC) to help identify more peptides and proteins (Swaney *et al*, 2010). LC-MS/MS analysis showed that 5179 proteins were identified with high confidence. Among them, 4716 proteins were identified in trypsin digestion and 4730 were identified in lysC digestion. The number of proteins identified in both datasets was 4267, consisting of 90.4% of trypsin digested samples and 90.2% of lysC digested samples, respectively (Fig 1D). The average sequence coverage of identified proteins in trypsin digestion was 29%, which was 2% higher than that in lysC digestion, as trypsin digestion generated more proteotypic, or easily detectable peptides for MS analysis (Fig S1A). The combination of two proteases digested dataset further improved the average sequence coverage to 36%, leaving significantly less proteins with low sequence coverage (Fig S1A). Though the application of trypsin and lysC digestion helped to identify more proteins with higher sequence coverage, it did not improve the identification of proteins with low molecular weight (LMW) (Fig S1B).

**Fig 1.**
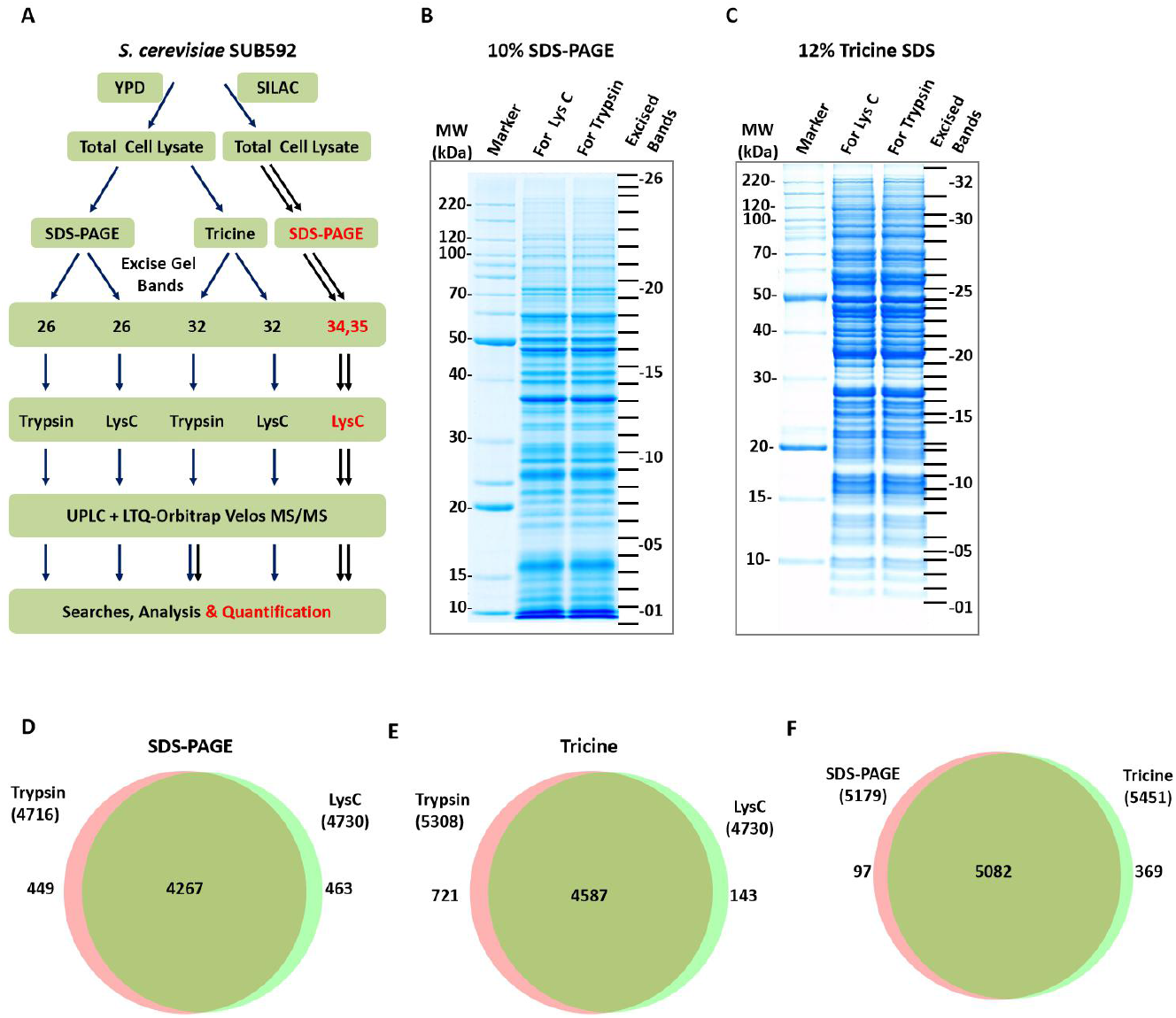
A nearly complete draft of the yeast proteome using MS-based proteomics. A, Three strategies used for the nearly complete coverage of yeast proteome. B, Sampling the yeast proteome by 10% SDS-PAGE and LC-MS/MS. C, Sampling the yeast proteome by 12% Tricine SDS-PAGE and LC-MS/MS. D, Venn diagram of proteins identified by SDS-PAGE by trypsin and lysC digestion. E, Venn diagram of proteins identified by Tricine SDS-PAGE by trypsin and lysC digestion. F, Venn diagram of proteins identified by SDS-PAGE and Tricine SDS-PAGE.

One way to increase the identification of LMW proteins in MS is to increase their resolution. Tricine gel has previously been shown to efficiently resolve LMW proteins with high resolution (Haider *et al*, 2012; Schagger, 2006). To identify more LMW proteins, we tested whether applying tricine gel can improve LMW proteins coverage (Fig 1C). Similar to the SDS-PAGE strategy, the samples resolved by tricine gel were also in-gel digested with trypsin or lysC and then analyzed by LC-MS/MS. The examination of MW distribution indeed indicated that the uniquely identified proteins from tricine gel were enriched in the region of LMW, and the number of identified proteins with MW<=10 kDa had improved by 31% (Fig S1C). The tricine gel runs resulted in a total of 5451 identified proteins (Fig 1E). Compared with the proteins identified from SDS-PAGE, 369 unique proteins were identified in tricine, increasing the total number of identified proteins to 5548 (Fig 1F). Compared to the published yeast proteome datasets (de Godoy *et al*., 2008; de Godoy *et al*., 2006; Ghaemmaghami *et al*., 2003; Huh *et al*., 2003; Nagaraj *et al*., 2012; Picotti *et al*., 2009; Picotti *et al*., 2013), our dataset is significantly larger, suggesting that protein identification has approached saturation using the current experimental conditions.

To further increase the number of identified yeast proteins, we reanalyzed our published proteome dataset derived from the same genetic background yeast strain cultured in synthetic complete (SC) medium for SILAC labeling (Li *et al*, 2019). The SILAC dataset increased protein identifications slightly from 5,548 to 5,610 (Fig S1D). Most of these additionally identified proteins were located in the LMW range (Fig S1E). Alteration of growth conditions did not significantly improve the number of identified proteins. This combined with the number of proteins identified from YPD experiments suggests that detection of proteins in all molecular weight ranges is likely approaching saturation. Therefore, the largest yeast proteome dataset to date is constructed with 5610 high-confidence gene products, covering 83.5% of yeast protein coding genes (Fig 2A, Supplementary table 2).

**Fig 2.**
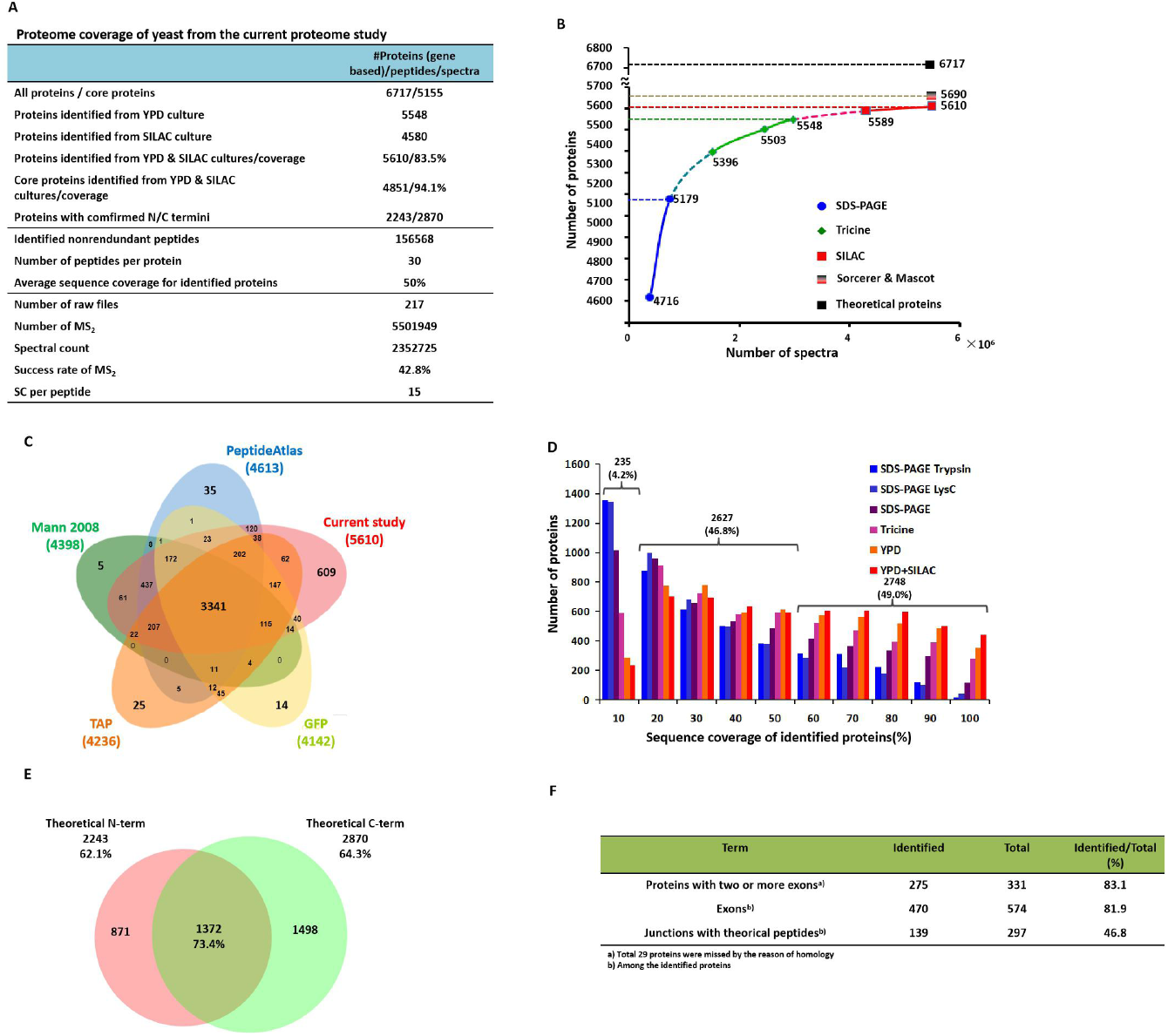
In-depth coverage of yeast proteome. A, Proteome coverage of current study. B, Number of identified proteins by the accumulated spectra from different approaches. C, Proteome coverage of current sutdy in comparison to previous studies. D, Sequence coverage of identified proteins by different experimental strategies. The number above the bracket represents the sum of the corresponding proteins. The percentage in the bracket represents the proportion of the corresponding proteins among all the proteins identified in this proteome. E, Venn diagram of the identified proteins having identified theoretical N- or C-terminal peptides in this proteome. The percentage below the number represents the average sequence coverage of the corresponding proteins. F, Identification of intron-containing gene products by this proteome.

Employing different experimental strategies not only increases the number of identified proteins, but also improves the accuracy of the identified proteins. Among the 5610 identified proteins, 97.1% matched at least two identified peptides, 99.2% matched at least one PSM with Xcorr>2 (Fig S2 A&B). The average number of identified peptides per protein reached up to 30, leading the average protein sequence coverage up to 50% (Fig 2D), which, to our best knowledge, is higher than the known proteomics studies to date (de Godoy *et al*., 2008; de Godoy *et al*., 2006; Nagaraj *et al*., 2012). It suggests the high reliability of our proteome dataset in protein identification. In SGD, yeast genes can be classified into three main categories: core, uncharacterized (including putative or hypothetical) and dubious genes. Among the 5155 core genes with annotated functions, 4851 were included in our dataset, reaching a coverage of 94.1% (Fig 2A&S2C, Supplementary table 2), indicating that the MS-based proteomics approach can reach near complete coverage for these core proteins. In addition, 71.4% of the uncharacterized genes and 27.4% dubious genes were identified in our dataset. All three catalogued gene groups were higher than the four previously published datasets (Fig S2C). Interestingly, our proteome provided support for the translation of 6 pseudogenes from 26 annotated ones in the reference yeast genome, in which YLL016W and YAL065C were uniquely identified in our study (Fig S2D). YLL016W was confirmed by the alignment of the spectra from large scale proteomics and synthesized peptides (Fig S2E).

Utilization of different experimental strategies helps to increase the number of identified proteins, however, as the accumulative spectra increases, less new proteins are identified (Fig 2B). MS-based experiments alone cannot efficiently improve the number of identified proteins, suggesting MS-based approaches have reached the upper limit of identification. In support of this, four published representative yeast datasets based on non-MS and MS techniques, consisted of Tandem Affinity Tag (TAP)-based dataset (Ghaemmaghami *et al*., 2003; Huh *et al*., 2003), Green Fluorescent Protein (GFP)-based dataset (Huh *et al*., 2003), PeptideAtlas dataset (Deutsch *et al*., 2008) and SILAC dataset published by Mann in 2008 (de Godoy *et al*., 2008), were selected to compare with our proteome dataset, we found very few novel proteins were identified based on these different datasets (Fig 2C). Most of the proteins uniquely in the other four datasets came from the GFP or TAP, which are not MS-based technologies and can play the role of complementing protein identifications. We further combined our dataset with these four datasets, which yielded a total of 5776 proteins by the aggregation of these five datasets, and 97.1% (5610) of these proteins were included by our dataset alone, suggesting the high coverage of our proteome dataset.

The high sequence coverage of the identified proteins help us confirm the annotation of the protein-coding ORFs in the current yeast genome, especially for the N-terminal and C-terminal ends of proteins. As protein termini may not generate proteotypic peptides long enough for mass spectrometric identification even using *in silico* digestion, here we defined the *in silico* digested peptide nearest to a protein terminus which could be identified by MS as the “theoretical terminus”, to represent protein terminus. As a result, 2,243 and 2,780 proteins had identified theoretical N-termini and C-termini, respectively, consisting of 40.0% and 49.6% of the identified proteins (Fig 2E). The average sequence coverage of these 2,243 and 2,780 proteins was 62.1% and 64.3%, respectively. A total of 1372 proteins had both identified theoretical N- and C-termini, with increased average sequence coverage up to 73.4%, which was significantly higher than that of all identified proteins in our proteome. We found that 799 and 1593 proteins had identified annotated N- and C-terminal peptides (Fig S3A), which provided the direct evidence of these proteins’ terminus annotation. Among the 779 proteins with annotated N-terminal peptide, 116 proteins had matched N-terminal peptide if the first amino acid residue in the N-terminus was removed, and 46 proteins had matched N-terminal peptide if the first two amino acid residues in the N-terminus were removed. Even still 8 proteins had matched N-terminal peptide after removing 5 amino acid residues (excluding targets amino acid of trypsin/lysC: lysine and arginine) from the N-terminus (Fig S3B). It indicates that a certain portion of yeast proteins has N-terminal cleavage sites of peptidase (Vogtle *et al*, 2009), which might regulate protein maturation, stabilization as well as function.

Another benefit of the high sequence coverage is reflected in the identification of intron-containing genes. In total we identified 275 of 331 (83.1%) annotated intron-containing gene products. Among these gene products, 470 exons were identified from the total 574, and 139 junctions were identified from the total 297, consisting of 81.9% and 46.8% respectively (Fig 2F). The amino acid sequence of junction peptide identified in YR111W-A was shown as an example in Fig S3C, further suggesting the high coverage of our proteomics data can provide direct evidence for the translation of gene splicing isoforms and facilitate the identification of splice sites.

### Characteristics of missing proteins in MS-based proteome study

Though our proteome dataset contains a total of 5610 proteins, there are still 1107 proteins missed based on SGD annotation. We performed a detailed analysis to uncover the possible reasons for the missing proteins.

Distribution of identified proteins based on MW as well as protein catalogue showed that proteins with LMW (≤20kDa) or belonging to uncharacterized or dubious gene products are mostly missed by our proteome dataset (Fig 3A). 840 of 1107 missing proteins were located in the LMW (≤20kDa) region (Supplementary Table S3). Proteins with LMW (≤20kDa) generate less peptides for MS-based proteomics to detect. Even when we applied tricine gel, which is optimized to identify small molecular weight proteins, still a large portion of proteins with LMW were left unidentified.

**Fig 3.**
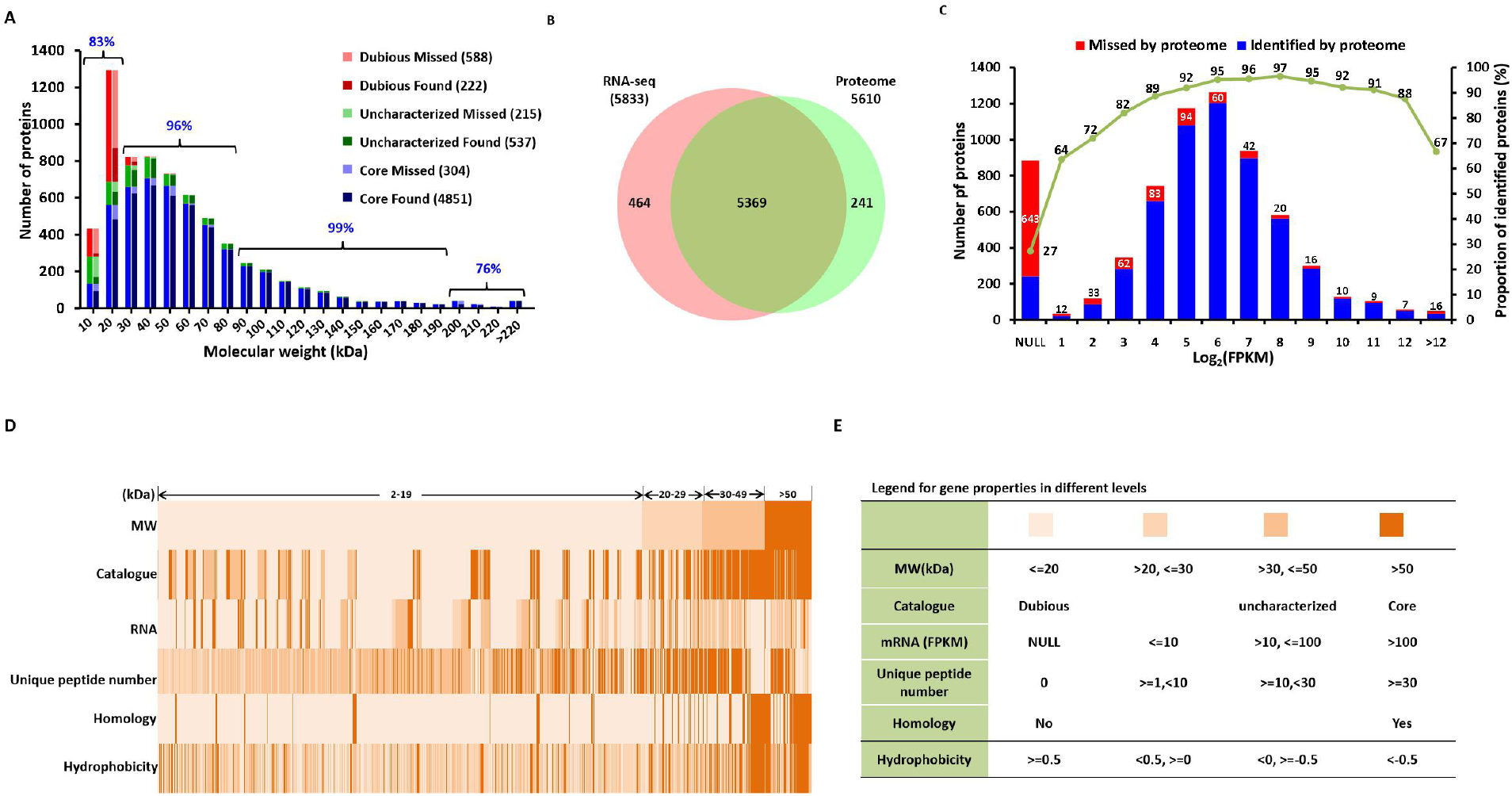
Characterization of missing proteins in our proteome. A, MW distribution of missed and identified proteins. The persentage of core proteins for the indicated MW range. B, Comparison of coverage by MS-based proteome and RNA-seq-based transcriptome (Li *et al*., 2019). C, Distribution of missed and identified proteins based on the mRNA abundance reflected by RPKM. The histogram represents the number of proteins identified (blue bars) or missed (red bars) by proteome in different bins of mRNA abundance. The green line represents the proportion of proteins identified by proteome in different bins of mRNA abundance. D, Distribution of 1107 missing proteins based on molecular weight, gene annotation, mRNA abundance, homology property, and protein physicochemical properties. Each column represents a missing protein. E, Legend for gene properties in different levels in D.

Compared to the nearly complete identification of core proteins, the identification of uncharacterized and dubious proteins were still low (71% and 27%) (Fig S2C), suggesting a large portion of these two categories proteins is still missing from our proteome dataset. Among 1107 missing proteins, a total of 803 proteins was uncharacterized or dubious proteins (Fig 3A, Supplementary table 3). Among, 723 proteins were also LMW proteins, consisting of 65.3% of the total missing proteins in our dataset.

The low identification of uncharacterized proteins as well as dubious proteins prompts us to explore whether the transcripts of these missing proteins are expressed or not with the assistance of RNA sequencing (RNA-seq). We compared our proteome dataset with our previously published RNA-seq dataset, which was performed in the same yeast strains under the same culture conditions (Li *et al*., 2019). The RNA-seq dataset contains 5,833 genes identified in total, representing an in-depth transcriptomics. A total of 5369 gene products were identified in common, occupying 95.7% and 92.0% of identified proteins and sequenced gene transcripts, respectively (Figure 3B). Among 1107 missing proteins, a total of 643 proteins were not detected in RNA-seq dataset (Fig 3C), including 525 uncharacterized or dubious proteins, suggesting under current growth conditions, a large portion of uncharacterized or dubious genes may not express. The following 464 missing proteins showed the normal distribution according to the RNA expression level, which is similar to the distribution of the identified proteins.

By comparing the proteomics data with protein MW and the RNA-seq dataset on a three-dimensional distribution, we found the missing proteins which were not detected by RNA-Seq are also of small MW (Fig S4). The union of missing proteins caused by LMW, uncharacterized and dubious protein categories and absence in RNA-seq dataset, is 986 proteins, consisting of 89.0% of the total missing proteins.

The remaining 121 missing proteins were all core proteins, with molecular weight ranging from 21 to 203 kDa. As for the identified core proteins, the coverage with MW≤20, 20-80, 80-190, >190kDa was 83.1%, 96%, 98.8% and 76.4% respectively (Fig 3A). It showed the lowest coverage of core proteins with MW>190kDa, even lower than the core proteins with MW≤20 kDa. This prompted us to analyze other physicochemical properties of these missing proteins. We found several of the missing proteins belonged to the retrotransposon protein group, which shared high sequence similarity. As peptides are the targets for sequencing in bottom-up shotgun proteomic strategies, proteins with highly conserved amino acid sequence will be mostly made up of non-unique peptides which are reported as a ‘protein homology group’ (Zhang *et al*, 2013). A parsimonious approach is to only choose one protein for each group, so the others are cataloged as missing proteins, though these proteins may have high sequence coverage. In fact, among the 1,107 missing proteins, 149 had at least one matched peptide, and 134 of the 149 proteins have more than 10% sequence similarity to identified proteins (Fig S5A). Most of these 134 proteins fall into three major protein groups, including retrotransposon, helicase, and ribosome (Fig S5B-D, Supplementary table 4). Therefore, proteins in these groups that are labelled as missing are primarily due to the high sequence similarity with the identified proteins, even though many of them have a high molecular weight (HMW) (Supplementary table 3). We found that 32 of 121 missing proteins in the core protein category belong to the highly homologous retrotransposon, helicase as well as ribosome groups. Thus, lack of unique peptides in HMW proteins remains a hurdle for complete coverage.

The hydrophobicity and number of proteotypic peptides have been proposed to account for the protein identification in MS (Amado *et al*, 1997; Krause *et al*, 1999). We found that the distribution of hydrophobicity or the number of proteotypic peptides were not significantly different between the identified proteins and the missing proteins (Fig S5 E&F). This indicates that our MS-based platform are robust enough to identify proteins regardless of their physicochemical parameters, further supporting the high sensitivity.

We also noticed that the distribution of the unidentified proteins are biased toward the ends of each chromosome (Fig S5G). More than 75% proteins localized near centromere were identified by either proteome or transcriptome, while only 50% proteins localized in chromosome ends were identified, which was extremely low in the chromosome extremities (~40%). This is likely due to the irregular repeated sequence of the telomeres in yeast, which differs from that of higher organisms including humans (Louis, 1995; Louis *et al*, 1994).

Hierarchical analysis for the integration of different protein characteristics showed that 1018 of 1107 missing proteins are caused by LMW, uncharacterized or dubious genes, absence in transcriptomics and sequence similarity (Fig 3 D&E, supplementary table S3). Among the 89 leftover uncharacteristic missing proteins, 45 did not generate enough proteotypic peptides for MS detection as predicted by peptideSieve, and 16 belonged to the enriched gene ontology (GO) catalogues associated with temporare expression, such as response to toxin, sexual sporulation or cell development (Fig S5H).

### Label-free quantification analysis shows the high correlation between the quantitative proteome and transcriptome

To correlate our proteomics dataset with gene expression, we quantitatively analyzed our label-free proteome based on peptide intensity. Because the abundance of different proteins could not be compared directly based on the intensity of all identified peptides due to the bias of peptide detectability by MS (Mallick *et al*, 2007), we designed a label-free workflow for combining quantitative results from different YPD experiments at the peptide level (Fig S6A). The peptides with abnormal intensity for each protein were eliminated due to the high sequence coverage in our proteomics dataset (Peptides identified from YML120C were shown as the example in Fig S6B), to further improve the accuracy of protein quantitation. Protein abundance was defined by the sum of the peptide intensities of each protein divided by their respective MW.

A total of 5056 proteins were quantified, comparable to the yeast unified protein abundance dataset, which combined 21 quantitative yeast proteome datasets (Ho *et al*., 2018). We found a large dynamic range of protein expression (Fig 4A), spanning approximately 6 orders of magnitude, which is 2 magnitudes larger than the mRNA abundance in the RNA-seq dataset (Li *et al*., 2019). This is consistent with what we find in human liver tissue (Chang *et al*, 2014a). Our quantitative proteome and the RNA-seq dataset had 4,923 gene products in common (Fig 4B). The Pearson correlation coefficient between the protein abundance and the mRNA abundance was 0.65 (Fig 4C), which is higher than our previous study based on quantitative SILAC method (Li *et al*., 2019), suggesting that the abundance of proteins is coupled with the abundance of mRNA (Marguerat *et al*, 2012). We also found that as the increasing of the number of quantitative peptides for each protein, the Pearson correlation of the intensity between transcriptome and proteome is also increased (Fig 4D), suggesting that increased depth of MS-based proteome in the future will improve quantitative accuracy and consistency with quantitative transcriptome, at least to some extent. Not only does our proteomics dataset correlate well with the transcriptomics dataset, it also correlates well with other published datasets that are generated with non-MS or MS based methods such as TAP (Ghaemmaghami *et al*., 2003) and GFP (Huh *et al*., 2003) (combined as TAP&GFP), as well as the quantitative SRM dataset (termed as SRM) (Picotti *et al*., 2013), with the respective Pearson correlation coefficients of 0.66 and 0.93 (Fig 4E, S6C). The high correlation with SRM dataset further suggests the high quantitative accuracy of our current proteomics dataset. As the quantitative information of SRM dataset is generated by the targeted comparison to the synthetic peptides with a known concentration (Picotti *et al*., 2013), which provide accurate relative quantification information for yeast proteins. Correlation coefficient between the transcriptome and TAP&GFP datasets was 0.51 (Fig 4F), which was lower than that with our proteomics dataset. Correlation coefficient between the transcriptome and the SRM dataset was, as expected, up to 0.83 (Fig S6D). Interestingly, it was lower than 0.93, which is the correlation coefficient between our proteomics dataset and the SRM dataset (Fig S6C). This suggests that our quantitative proteomics dataset better reflects the relative gene expression pattern, compared to the quantitative transcriptome dataset. It is likely due to the post-transcriptional regulation via control over translation and/or degradation rates of specific proteins within the cell (Tchourine *et al*, 2014).

**Fig 4.**
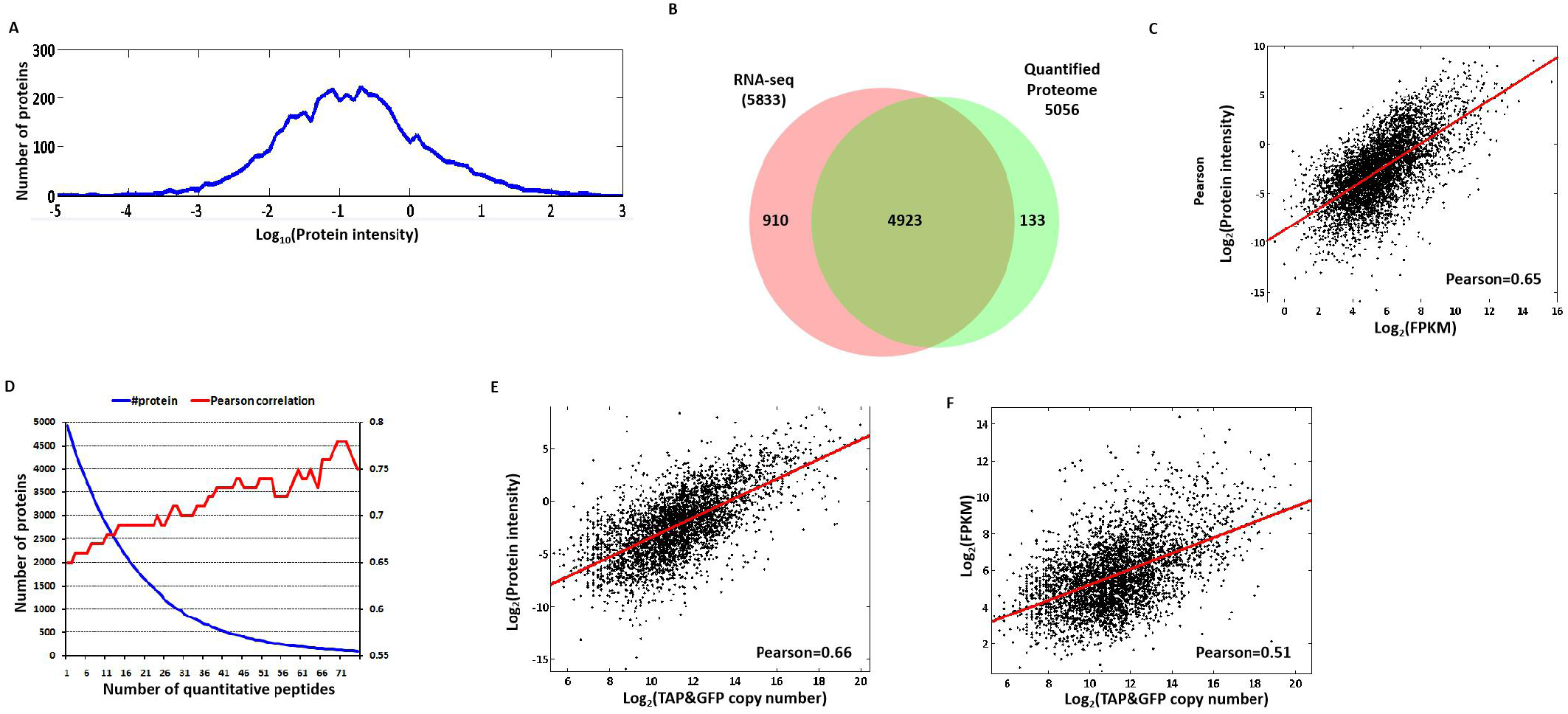
High correlation of our quantified proteome with trancriptome. A, Dynamic range of protein abundance. B, Comparison of the coverage of quantified proteome and RNA-seq-based transcriptome(Li *et al*., 2019). C, Correlations between quantified proteome and transcriptome (Li *et al*., 2019). The *x*-axis represents the log_2_ FPKM, and the *y*-axis represents the log_2_ protein intensity. D, The curve of the number of quantitative peptides for a protein and the pearson correlation of the intensity between proteome and transcriptome. The *x*-axis represents the number of quantitative peptides for each protein. The left *y*-axis represents the number of proteins corresponding to the number of quantitative peptides, and the right *y*-axis represents the pearson correlation of the intensity between proteome and transcriptome for these proteins. E, Correlations between our quantified proteome and TAP&GFP datasets (Ghaemmaghami *et al*., 2003; Huh *et al*., 2003). The *x*-axis represents the log_2_ protein copy number in TAP&GFP datasets, and the *y*-axis represents the log_2_ protein intensity in our quantitative proteome. F, Correlations between TAP&GFP datasets (Ghaemmaghami *et al*., 2003; Huh *et al*., 2003) and transcriptome (Li *et al*., 2019). The *x*-axis represents the log_2_ protein copy number in TAP&GFP datasets, and the *y*-axis represents the log_2_ FPKM.

To further quantitatively compare our proteomics dataset with the TAP and GFP datasets, we transformed our protein intensity into the copy number using the SRM dataset as a ruler (see method) (Supplementary table 2) (Picotti *et al*., 2013). The dynamic range of protein copy number in our dataset was two magnitudes larger than that given by TAP and GFP construct expression, extending mainly in the direction of low protein abundance (Fig S6E&F). Our proteomic dataset identified 241 and 609 unique proteins not found by RNAseq (Fig 3B) and the four other published datasets (Fig 2C), respectively. Additionally, we also showed a biased distribution in the low expression region, both in protein and RNA level (Fig S7). Hence, identification of low-abundance proteins drives the improvement towards complete coverage in our proteomic dataset, and reflects the depth of our MS-based pipeline.

### Functional pathway profiling by the high coverage quantitative proteome

Our quantitative proteome dataset analysis provides insight into the protein expression pattern of yeast under the log phase growth conditions (Fig 5A). The core proteins have globally higher abundance than the uncharacterized proteins and the products of dubious genes (Fig S8), which further suggests that these core proteins are essential to yeast. This is consistent with what we found in our previous SILAC dataset (Li *et al*., 2019).

**Fig5.**
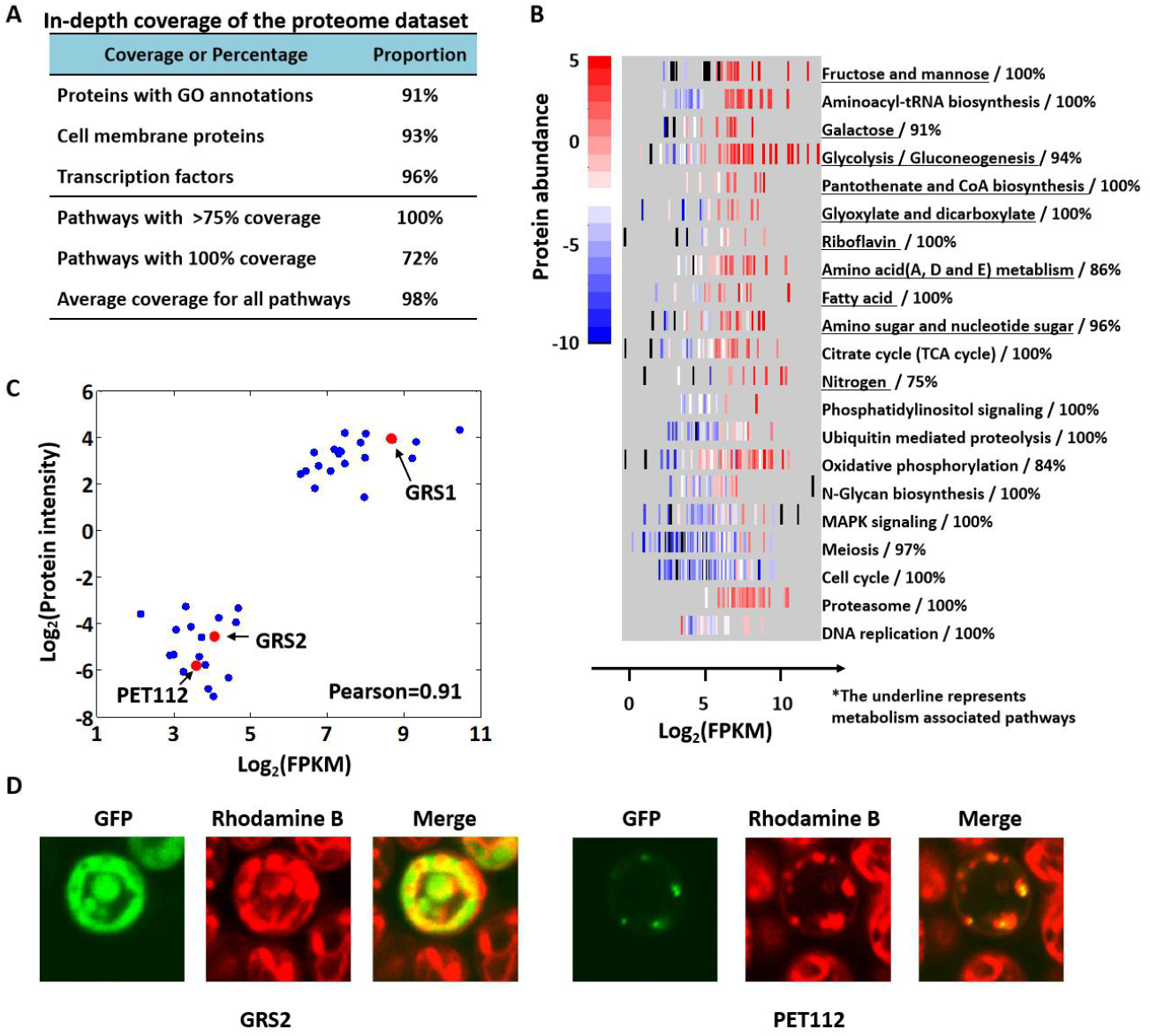
Functional protein-coding genes and pathways profiling based on our quantitative proteome. A, Protein coverage of the different biological pathways. B, 21 KEGG pathways with high correlations between transcriptome and quantified proteome. Top 21 pathways enriched by the quantitative proteins were selected, and ranked by the correlation of transcriptome and quantified proteome from high to low. Different colors represent different abundance of proteins. Blank refers to the proteins that cannot be quantified in proteome. The percentage on the right represents the proteome coverage for each pathway. C, Two groups of aminoacyl-tRNA biosynthesis enzymes based on their protein/RNA abundance. The correlation between transcriptome and proteome for these genes was analyzed. GRS family was highlighted in red. D, Visualization of the mitochondrial localization of the C-terminally GFP-tagged GRS2 and PET112 by confocal microscopy. The three images show the same group of cells visualized by fluorescence using the GFP (GFP), or the rhodamine B hexyl ester (Rhodamine B) channels, or an overlay of the GFP signal to Rhodamine B signal (Merge).

All intracellular components attain high identification coverage (>93%), except for the extracellular region and cell wall (72.6% and 74.8%, respectively). Even membrane proteins, which can be difficult to extract, digest, and detect in such experiments, also attain 93.4% coverage (Fig S9A). Besides that, 96% of transcription factors and 91% of all proteins with GO slim annotations were covered in our proteomics dataset, providing additional evidence that most of the annotated functional protein-coding genes are expressed in yeast cells under log-phase growth conditions.

Our proteomic dataset covers almost all proteins essential for yeast survival as supported by pathway analysis. The coverage of all proteins in the KEGG pathway were above 75%, with 72% of pathways having all their proteins completely covered (Supplementary table 5); the average coverage of KEGG pathway annotated proteins is 98% (Fig 5A). One of the most active pathways, mitosis, is chosen for detailed analysis. Mitosis associated proteins are cataloged into five subgroups (midbody, centrosome, kinetochore, telomere and spindle) based on the microkit 4.0 (Ren *et al*, 2010) and SGD annotations. More than 97% of all five subgroups of their member proteins were uniquely identified (Fig S9B, missing proteins are listed in Supplementary table 6).

Combining mRNA and protein abundance to the proteins assigned in each KEGG pathway further uncovered the expression patterns of different functional modules under current growth conditions. Fig 5B presented proteins in representative pathways with mRNA and protein abundance; pathways were ranked by the correlation coefficient between the transcriptome and the proteome from high to low. This confirms that (1) the correlation of protein to mRNA is higher not only for individual genes, but also extend to the well-established pathways; (2) protein encoding genes in the concerted metabolic pathways have high correlation with their transcript levels, suggesting that the transcriptional control is a primary means of regulating the abundance of these proteins; (3) proteins involved in meiosis and cell cycle have relatively low correlation with their transcript abundance, possibly due to stringent regulation of checkpoint controls where protein expression might lag behind mRNA changes such as multiple post-translational modification to achieve necessary changes in function.

Subcellular localization of proteins is an important aspect of gene annotation, which relates to its cellular function. It has been previously shown that protein abundance and localization is regulated together (Torres *et al*, 2016). Here our quantitative proteome dataset with accurate protein abundance information provides a proteome-wide view of protein expression pattern, including protein subcellular localization. Using proteins in the aminoacyl-tRNA biosynthesis pathway as examples, we show that correlation of mRNA and protein abundance of this pathway is 0.91 (Fig 5C). All 39 proteins can be classified in 2 groups based on their mRNA abundance and protein abundance. Among the 21 high abundance proteins, 13 were annotated to localize in cytoplasm; 17 of the 18 low abundance proteins were annotated to localize in mitochondria. The one remaining low abundant protein (GRS2) is currently left unannotated in the SGD is probably localized in mitochondria. Confocal microscopy analysis confirms that GRS2 is indeed located in mitochondria (Fig 5D).

## Discussion

In MS-based shotgun proteomics, a longstanding challenge is to identify the entire set of proteins that are complementary expressed by a genome, cell or tissue type (de Godoy *et al*., 2008; Kim *et al*., 2014; Mergner *et al*., 2020; Nagaraj *et al*., 2012; Picotti *et al*., 2009; Wilhelm *et al*., 2014). Sophisticated sample preparation and separation, high sequencing speed and sensitivity have significantly improved the protein identification in many species (Domon & Aebersold, 2006; Kumar & Mann, 2009; Shevchenko *et al*., 1996b; Washburn *et al*., 2001). Here, we take full advantage of the molecular size based separation that is enabled by high resolution SDS-PAGE, optimized LC gradient (Xu *et al*, 2009) and high resolution Orbitrap Velos MS (Li *et al*, 2012) to generate full coverage of yeast proteome. We have identified 5610 proteins in total, with their abundances spanning across nearly six orders of magnitude (Fig 4A). 94.1% of the theoretical core proteome has been identified (4851). 71% and 22% uncharacterized and dubious gene products (537 and 222) are identified (Fig S2C). The remaining unidentified proteins are due to LMW, absence in transcription or high sequence similarity (Fig 3). This is considerably higher than the previous comprehensive proteomics studies of yeast (de Godoy *et al*., 2008; Deutsch *et al*., 2008; Ghaemmaghami *et al*., 2003; Huh *et al*., 2003). We also demonstrate that our high quality dataset can facilitate gene annotation as well as gene expression pattern in defined growth conditions.

We have utilized label-free as well as SILAC strategies under different growth conditions to generate spectra using our MS platform. We find that past a certain point there is a negative correlation between increasing spectra number and additional proteins identified (Fig 2B), suggesting the approach of a saturation point. SDS-PAGE gel-based label-free method identifies 5179 proteins. Combining SDS-PAGE gel- and tricine gel-based label-free methods increases identification to 5548 proteins. Combining all label-free and SILAC methods brings an increase of only 62 proteins and a total of 5610. This indicates that more large-scale MS-based experiments cannot efficiently increase the number of identified proteins, even though different strategies of digestion and separation are used. As for the bioinformatics analysis, another search engine, Mascot (Perkins *et al*, 1999), only added 80 more proteins with low quality (data not shown), hence these proteins are not included in our proteome dataset. These analyses suggest that our proteome dataset has reached the limit for the yeast proteome, at least for the MS-based methods.

Based on 6717 annotated yeast ORFs in SGD database, 1107 proteins are missing in our proteome dataset. We comprehensively analyze the characteristics of these 1107 missing proteins from protein physicochemical properties to protein expression, which may provide new clues for further improving proteomics study. We find that LMW, absence in transcriptome dataset, uncharacterized and dubious genes, and high sequence similarity account for almost all of the missing proteins annotated in SGD. For example, among the 304 core proteins missed by our proteome dataset, 117 are proteins with MW<=20 kDa, 104 are highly homologous with identified proteins, and 118 are missed by RNA-seq dataset. The combination of these three catalogues (LMW, high sequence similarity and absence in transcriptome) proteins are 215, leaving 89 proteins as part of the denominator. In this way, the fixed proteome coverage of core proteins reaches 4274/(4274+89)=98.0%, indicating that MS-based proteomics technology achieve near complete coverage for basic ORFs (Fig 3D&E). These results further confirm the near complete coverage of our proteome dataset.

Integrative analysis of our proteomics data and in-depth RNA-seq data not only help to figure out the reason for missing proteins, but also provided insights into the global proteomics dynamics and function of metabolic and cellular regulatory networks in yeast. Protein abundance of our proteomics data spans approximately 6 orders of magnitude, one magnitude larger than that in the previous 21 quantitative yeast proteome datasets (Ho *et al*., 2018) and 2 magnitudes larger than the mRNA abundance (Fig 4A, (Li *et al*., 2019)), suggesting the high sensitivity of our MS platform.

Our nearly complete proteome dataset can also be used to validate and revise yeast genome annotation. It can help to characterize protein N- or C-terminal sequence, and to provide expression evidence of pseudogenes. Moreover, based on the accurate protein abundance information, it can also provide reliable information about protein localization in cells (Fig 5C&D). These results suggest that our proteome dataset would be a useful blueprint for yeast proteogenomics study, to further optimize yeast genome annotation.

In conclusion, we provide the largest yeast proteome dataset so far based on MS technology, and highlight the characteristics and some of many uses that can be applied of this resource. These advances, combined with the fast multi-omics studies, will make the complete yeast proteome map possible for the foreseeable future.

## Materials and Methods

### Yeast Strains, medium and cultured protocols

Yeast strains used in this study were described in supplementary table 1. Yeast strains SUB592 were used in this study for yeast proteomic study. MHY500 was used to study localization of GRS2 and PET112.

To investigate the localization of GRS2 and PET112 proteins in yeast cells, we generated the plasmid expressing GFP-GRS2 or GFP-PET112 fusion proteins. The DNA fragments of Grs2 and Pet112 were amplified from JMP024 by colony PCR (Grs2-F: 5’-GGGGTACCATGCCGTTAATGTCCAATTCGG-3’; Grs2-R: 5’-TAGCGGCCGCATATCTTAACAGGCGACAGTCC; Pet112-F: GGGGTACCATGTTGCGGCTTGCACGT; Pet112-R: TAGCGGCCGCACCATTGAATATTTAAGATCTC-3’). The plasmids were made by inserting Grs2 or Pet112 into the pYES2-GFP vector (a gift from Dr. Matther J Higgins) using *Kpn*I and *Not*I sites, resulting in plasmids pYES2-GRS2-GFP and pYES2-PET112-GFP, respectively (Supplemental table 1). In these plasmids, GRS2 or PET112 was tagged at the carboxy terminal end with green fluorescent protein (GFP), under the control of the inducible GAL1 promoter. Then plasmids pYES2-GRS2-GFP and pYES2-PET112-GFP were transferred to strain MYH500 (Swanson *et al*, 2001), screened by SC medium without uracil to generate the strain PX001 and PX002, respectively. In addition, transformations were carried out according to the standard LiOAc method (Gietz & Woods, 2002).

In general, yeast strains were grown at 30°C in YPD medium (1% yeast extract, 2% Bacto-peptone, and 2% dextrose) and harvested at A_600_ of 1.5 unless indicated. The SC medium (0.67% yeast nitrogen base, 2% glucose, and supplemented with the appropriate amino acids) was used to generate yeast strains PX001 and PX002.

### Sample preparation for yeast S. cerevisiae and mass spectrometric analysis

The yeast strain *S. cerevisiae* SUB 592 was grown at 30°C in YPD medium, and harvested at the mid exponential phase. Cells were lysed in a 1.5 mL centrifuge tube with denaturing lysis buffer (8 M urea, 50 mM NH_4_HCO_3_, 10 mM IAA) and 0.5 mm glass beads (Biospec Products Inc., Bartlesville, OK). Protein concentration of yeast lysate was measured by a Coomassie stained SDS gel(Xu *et al*., 2009). The certain amount of TCL was separated through SDS-PAGE and Tricine gel and sliced into 26-35 fractions based on molecular weight markers and digested with trypsin or Lys C, respectively. After digestion overnight, the peptides were extracted in the extraction buffer (5%FA+45%ACN) and ACN, and finally dried with the vacuum dryer (Labco, CENTRIVAP).

Peptides were analyzed using a LC-MS/MS platform of hybrid LTQ-Orbitrap Velos mass spectrometer (Thermo Fisher Scientific, San Jose, CA, USA) equipped with a Waters nanoACQUITY ultra performance liquid chromatography (UPLC) system (Waters, Milford, MA, USA) as described previously (Li *et al*., 2019).

### Database searching for protein identification

Database searching was operated as described previously (Li *et al*., 2019). Briefly, all raw files were converted into mzXML using Trans-Proteomic Pipeline (version4.5.2) (Xu *et al*., 2009), and searched by the Sequest-Sorcerer algorithm (version 4.0.4 build, Sage-N Research, Inc, Sage-N-Research, Inc., San Jose, CA, USA) (Pedrioli, 2010) against the combined target-decoy proteins from *Saccharomyces* genome database (version released in 2011.02, 6717 entries http://www.yeastgenome.org/) along with 112 common contaminants (ftp.thegpm.org/fasta/cRAP).

The same parameters were employed for Mascot (version, 2.3.0) search (Chang *et al*., 2014a). The application of additional search engine can improve the identification coverage, but induce more false positive results (Cox & Mann, 2008). So we only adopted the results from the sorcerer software.

We also constructed a sequence database with different splices for the proteins with more than two exons, and searched it with the sorcerer software. As a result, no positive peptides were found.

### Protein quantitation

Label-free quantitation was operated as described previously (Li *et al*., 2019). The area under the extracted ion chromatograms (XICs) for each full digestion peptide in the YPD sample was calculated using SILVER (Chang *et al*, 2014b). As shown in supplementary fig 6, the intensity of a peptide was firstly normalized by the median of all peptide intensities in the corresponding sample, then the geometric mean of the intensities from four samples was calculated as the final intensity for each peptide. The mean and standard intensity of the unique peptides from the same protein was calculated. The peptides with intensity out of mean±2sd were removed as isolated points. The sum of the remaining peptides was divided by the protein MW as the final intensity of each protein.

### Bioinformatics analysis of identified peptide and proteins

Protein information, including gene symbol, chromosome loci, gene model and modifications, was mainly generated from SGD annotations. Four published datasets, Tandem Affinity Tag (TAP) (Ghaemmaghami *et al*., 2003), Green Fluorescent Protein (GFP) (Huh *et al*., 2003), PeptideAtlas (Deutsch *et al*., 2008) and Mann 2008 (de Godoy *et al*., 2008), were selected to compare with our proteome dataset. According to the SGD annotations, all proteins were classified into three catalogs including “Core”, “uncharacterized (including Putative or Hypothetical)” and “Dubious”. Core proteins represent the verified ORFs or the uncharacterized ORFs with essential function. “Put or Hypo” proteins represent the putative or hypothetical uncharacterized ORFs. “Dubious” proteins represent the dubious ORFs. Protein molecular weight and hydrophobicity were calculated using ProPAS (Wu & Zhu, 2012). Proteotypic peptides were predicted by PeptideSieve with threshold scores larger than 80 (Mallick *et al*., 2007). GO enrichment analysis was achieved by DAVID (http://david.abcc.ncifcrf.gov/) (Huang *et al*, 2009), and GO-slim information was generated from online tool GOTermMapper (http://go.princeton.edu/cgi-bin/GOTermMapper). Pathway information came from the database Kyoto Encyclopedia of Genes and Genomes (KEGG, http://www.genome.jp/kegg/) (Kanehisa, 2002). Mitosis annotations were generated from database MiCroKiTS 3.0 (http://microkit.biocuckoo.org/) (Ren *et al*., 2010). Venn was drawn by the online tool jvenn (http://bioinfo.genotoul.fr/jvenn/example.html) (Bardou *et al*, 2014). The figure of the cell structure was drawn using business software SmartDraw (http://www.smartdraw.com/).

### MS analysis of synthesized peptides for validation of pseudogenes

Peptides for validation of pseudogenes were synthesized and roughly purified (Shanghai Leon Chemical Ltd., Shanghai, China). The peptides (0.1-1pmol) were dissolved in ddH2O and desalted with homemade Stage-Tip (Zhai *et al*, 2013) and analyzed with LC-MS/MS as described above.

### Confocal fluorescence microscopy

The strain PX001 and PX002 were grown in SC medium to early-exponential phase (A_600_=0.7) and then washed three times by SC medium without glucose. Then GFP-GRS2 and GFP-PET112 fusion proteins were induced for 3 hr by addition of 2 % galactose. For staining of mitochondria in living cells, cultures of exponentially growing PX001 and PX002 were resuspended in 10 mM HEPES (Ph 7.4), 5% (w/v) glucose, 100 nM rhodamine B hexyl ester and incubated at room temperature for 30min. Cells were visualized with a Zeiss LSM510 META confocal fluorescence microscope with 40x objective. GFP was excited with a 488 nm laser, and its emission was collected at 509 nm, while rhodamine B hexyl ester was excited with a 555 nm laser and its excitation collected at 577 nm.

### Data availability

All the proteome raw and meta data was uploaded on proteomeXchange (http://www.proteomexchange.org/) with ID PXD001928.

## Acknowledgements

We are indebted to Drs. Fuchu He, Junmin Peng and Ning Li for support in the early stage of this project. We are grateful to Simin He, Hao Chi, Lanlan Li, Hui Jiang and Baoqing Ding for gracious gifts of their reagents, discussion, critical reading and editing. This work was funded by the State Key Development Program for Basic Research of China (2017YFA0505100, 2017YFA0505000 & 2016YFA0501300), the National Natural Science Foundation of China (31700723, 31670834, 31870824 & 91839302), the Innovation Foundation of Medicine (19SWAQ17, AWS17J008 & BWS17J032, 16CXZ027), National Megaprojects for Key Infectious Diseases (2018ZX10302302), Research Unit of Proteomics & Research and Development of New Drug of Chinese Academy of Medical Sciences (2019RU006), Guangzhou Science and Technology Innovation & Development Project (201802020016), the Unilevel 21st Century Toxicity Program (MA-2018-02170N), and the Foundation of State Key Lab of Proteomics (SKLP-K201704 & SKLP-K201901).

## Author contributions

YG and LP conceived the project. YG, LP and DD performed the experiments. CZ, ED, YL, PC, PX and LC analyzed the data. YG and PX wrote the manuscript with input from all authors. JW and PX oversaw the project.

## Supplemental figures

**Fig. S1.**
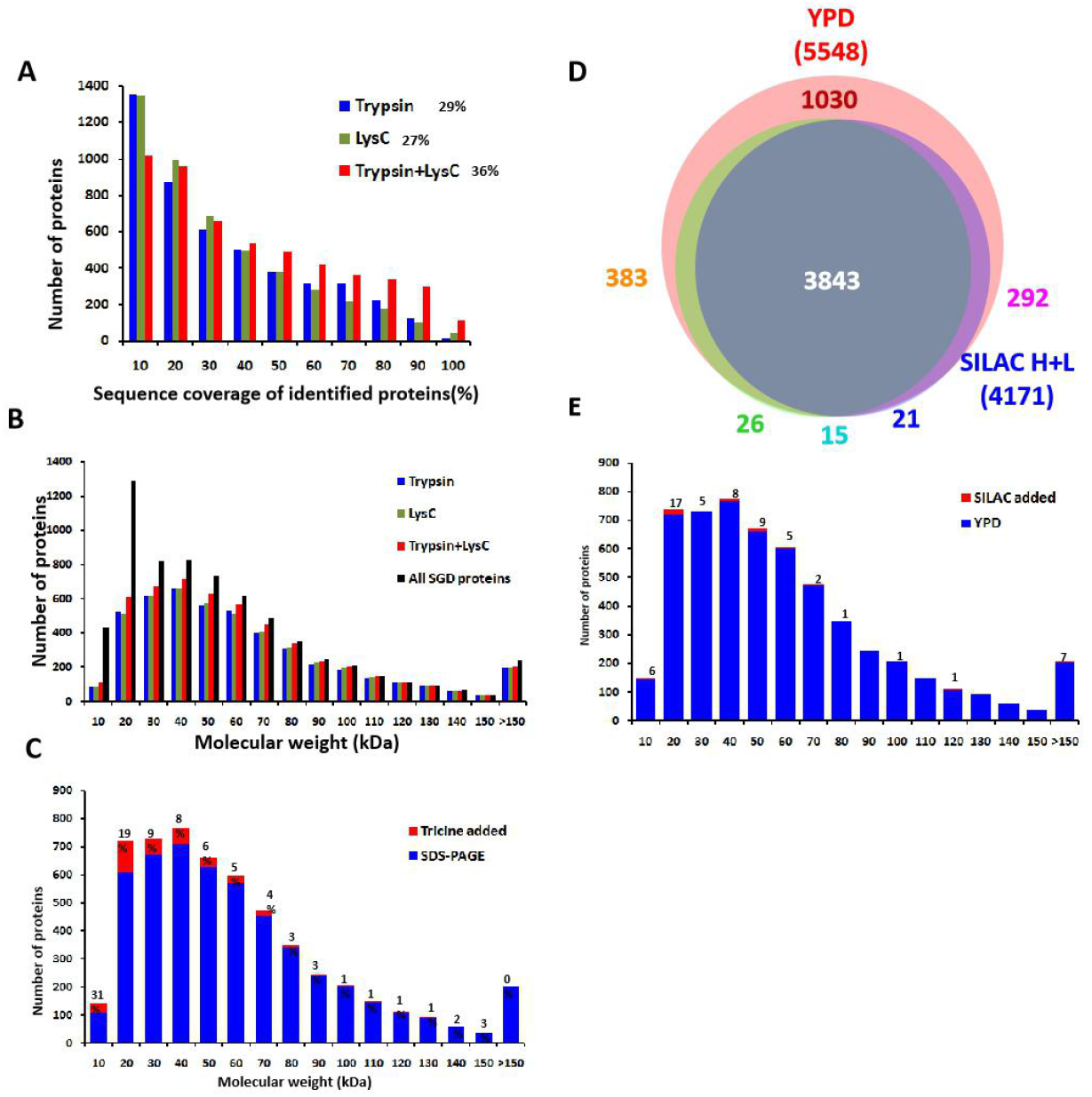
Contribution of different experimental strategies for deep proteome coverage. A, Distribution of the sequence coverage of identified proteins by trypsin and lys C in SDS-PAGE method. The number on the left of the legend represents the average sequence coverage of the corresponding identified proteins. B, MW distribution of theoretical and identified proteins by trypsin and lys C in SDS-PAGE method. C, MW distribution of added proteins identified by Tricine SDS-PAGE based on the result of SDS-PAGE. Percentage represents the proportion of identified proteins added by the Tricine SDS-PAGE. D, Venn diagram of identified proteins by YPD and SILAC (Li *et al*., 2019) medium. E, MW distribution of added proteins identified by SILAC dataset based on the result of YPD dataset. Number represents the number of identified proteins added by SILAC dataset.

**Fig. S2.**
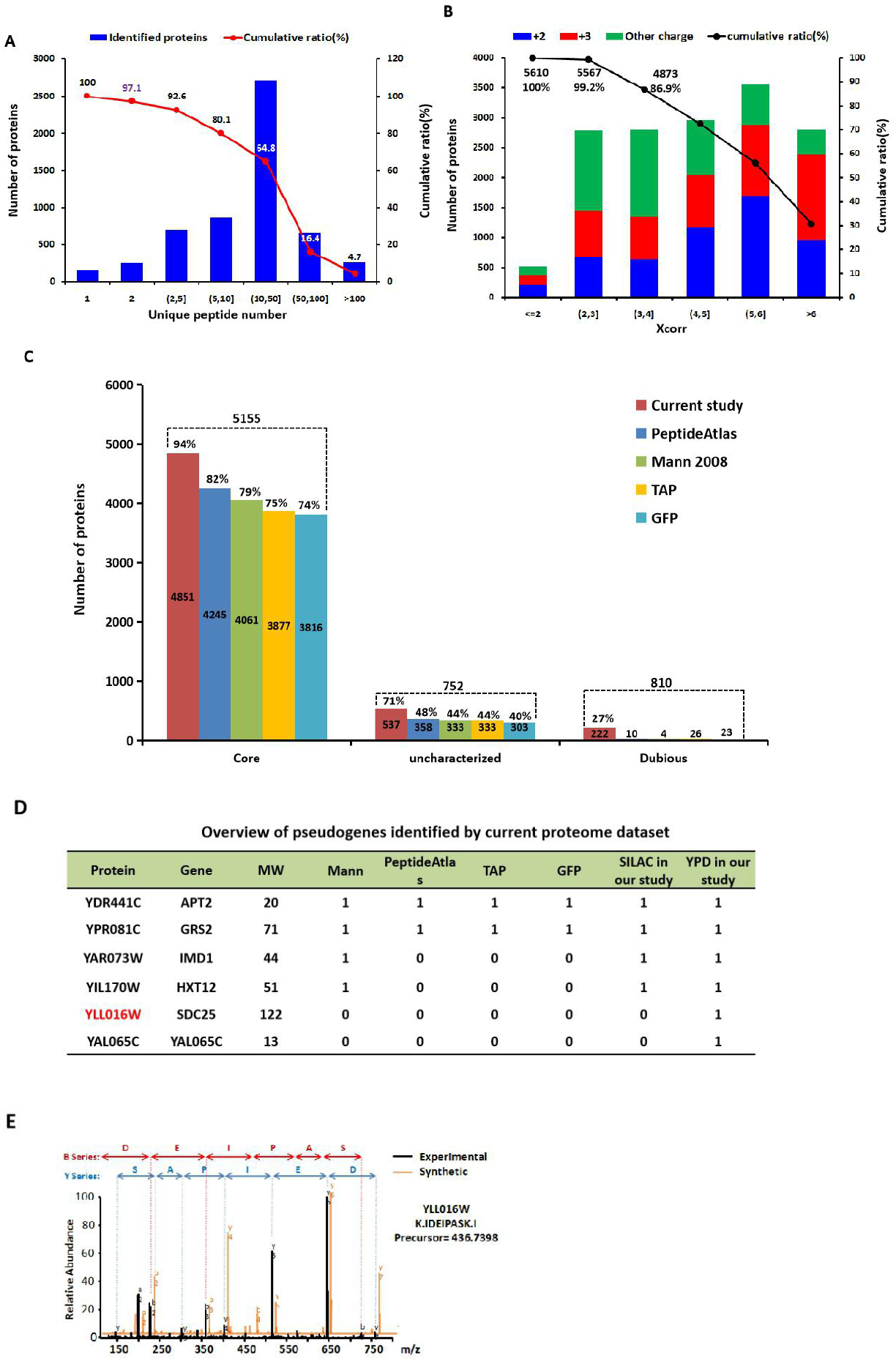
High coverage of different protein categories proteins by our proteome dataset. A, Number of unique peptides in identified protein. The number on the left *y*-axis represents the sum of proteins among each bin of peptide number. The percentage on the right *y*-axis represents the cumulative ratio of proteins with peptides greater than or equal to each bin. B, Distribution of Xcorr value assigned for identified proteins. The number on the left *y*-axis represents the sum of proteins among each bin of Xcorr value. The percentage on the right *y*-axis represents the cumulative ratio of proteins with Xcorr value greater than or equal to each bin. C, Comparison of proteome coverage of MS-based proteomic strategies from this study with four datasets of Mann 2008, Peptide Atlas, GFP- and TAP-tagging methods among the categories of core, uncharacterized (putative or hypothetical), and dubious proteins. Number above the dotted line represents the sum of each catalogue. Percentage above the bar represents the coverage of each dataset for the corresponding catalogue. D, Overview of the pseudogenes identified by our proteome dataset. Pseudo genes YLL016W was selected for validation. E, Comparison and validation of the MS2 spectra of the identified peptide generated from the pseudogene YLL016W in large scale proteomics with that of synthesized peptide.

**Fig S3.**
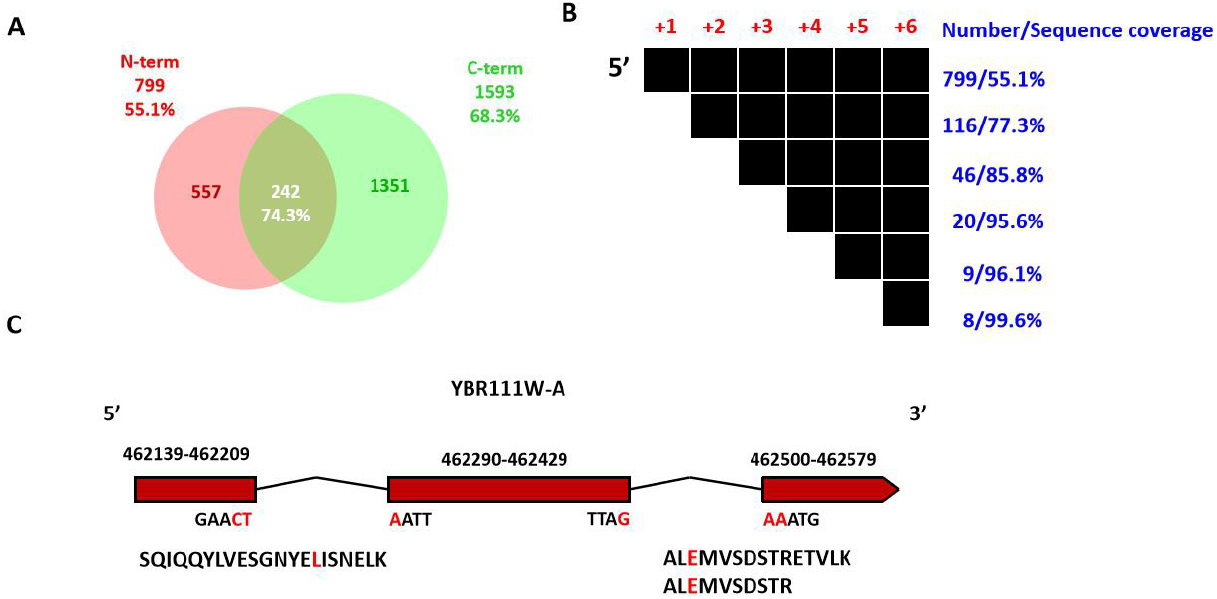
Validation of protein N- and C-termini sequence and splicing site based on identified spectra by our MS platform. A, Venn diagram of the identified proteins having annotated N- or C-terminal peptides identification in our proteome. The percentage below the number represents the average sequence coverage of the corresponding proteins. B, Number of proteins with identified peptides covering different sites in the N-termini. Each black block represents an amino acid covered by an identified peptide. The top line represent the proteins with identified peptides which have the whole exact N-termini in the corresponding proteins. Among the proteins belonging to the top line, if a protein owns identified peptides with N-termini located on the second amino acid of the protein N-termini, it would be cataloged into the second line. The same rule was applies to the other four lines. Percentage represents the average sequence coverage of the proteins in the corresponding line. C, Identification of the ‘junction’ peptides in YBR111W-A. The nucleotides refers to the sequence of junction after splicing, corresponding to below peptide identified in this study.

**Fig S4.**
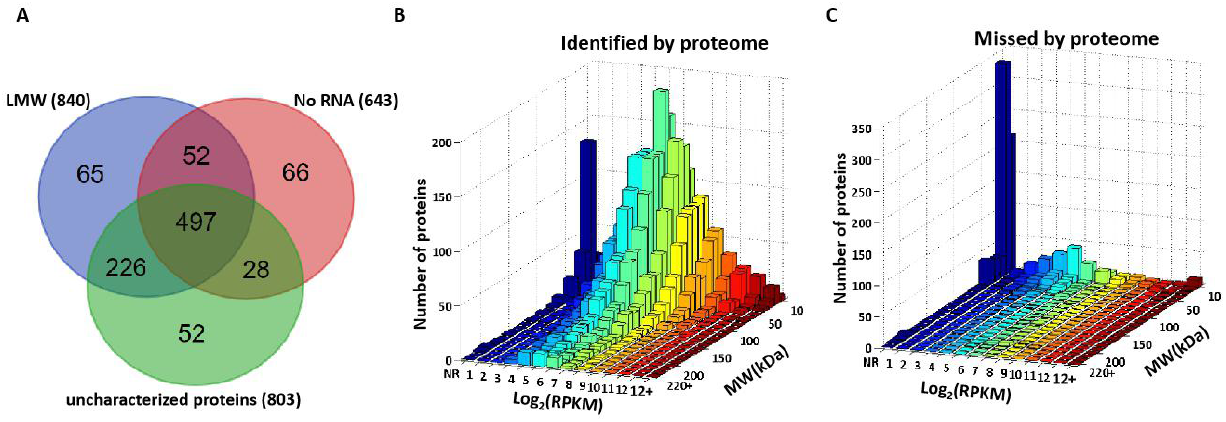
Overlapping of missing proteins belonging to LMW, no RNA expression and uncharacterized proteins. A, Venn diagram of the missing proteins belonging to LMW, no RNA expression and uncharacterized proteins. B&C, 3-Dimensional distribution of identified (B) and missing (C) proteins vs their theoretical MW and mRNA abundance. NR, not detected in RNA-seq dataset.

**Fig. S5.**
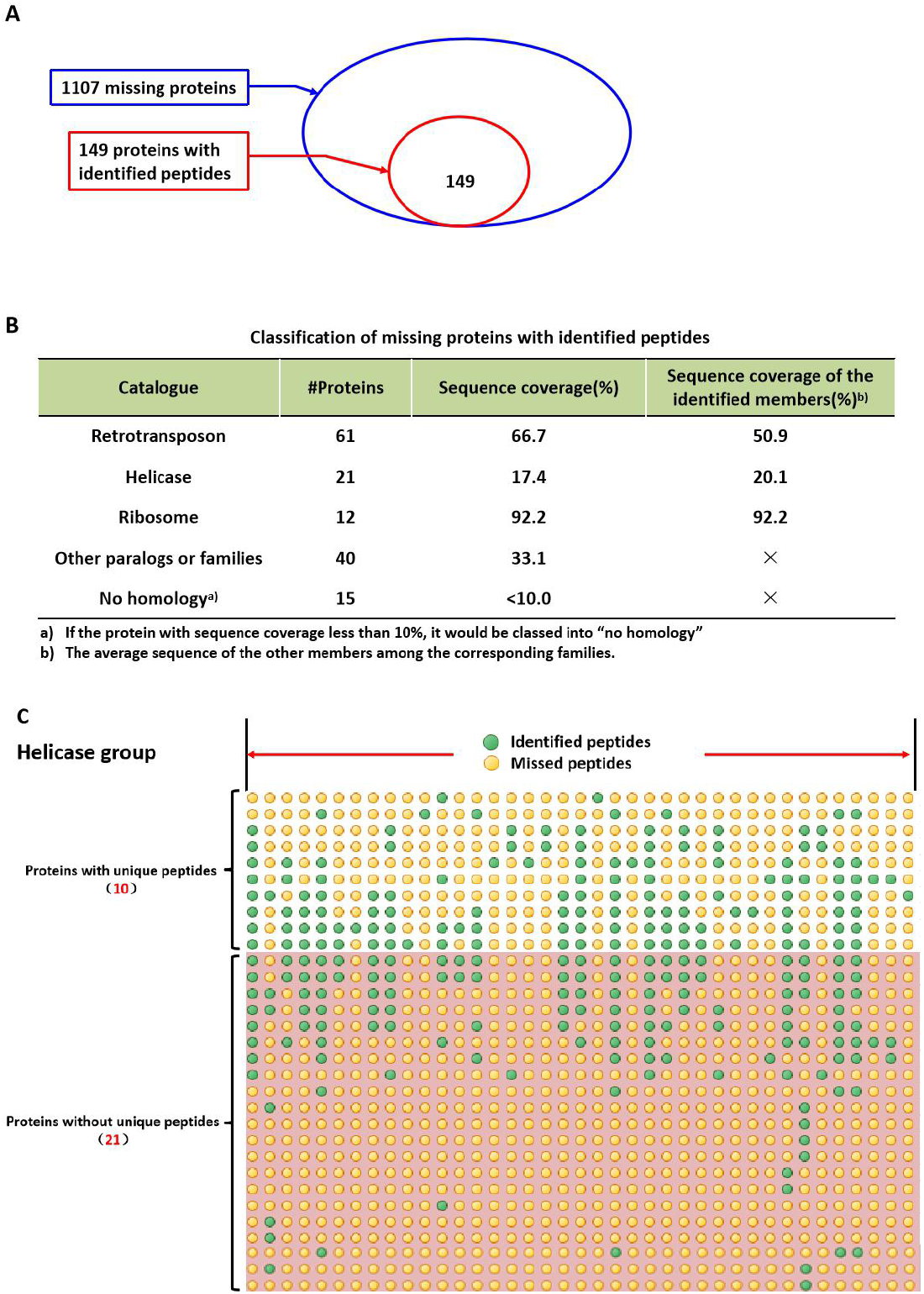
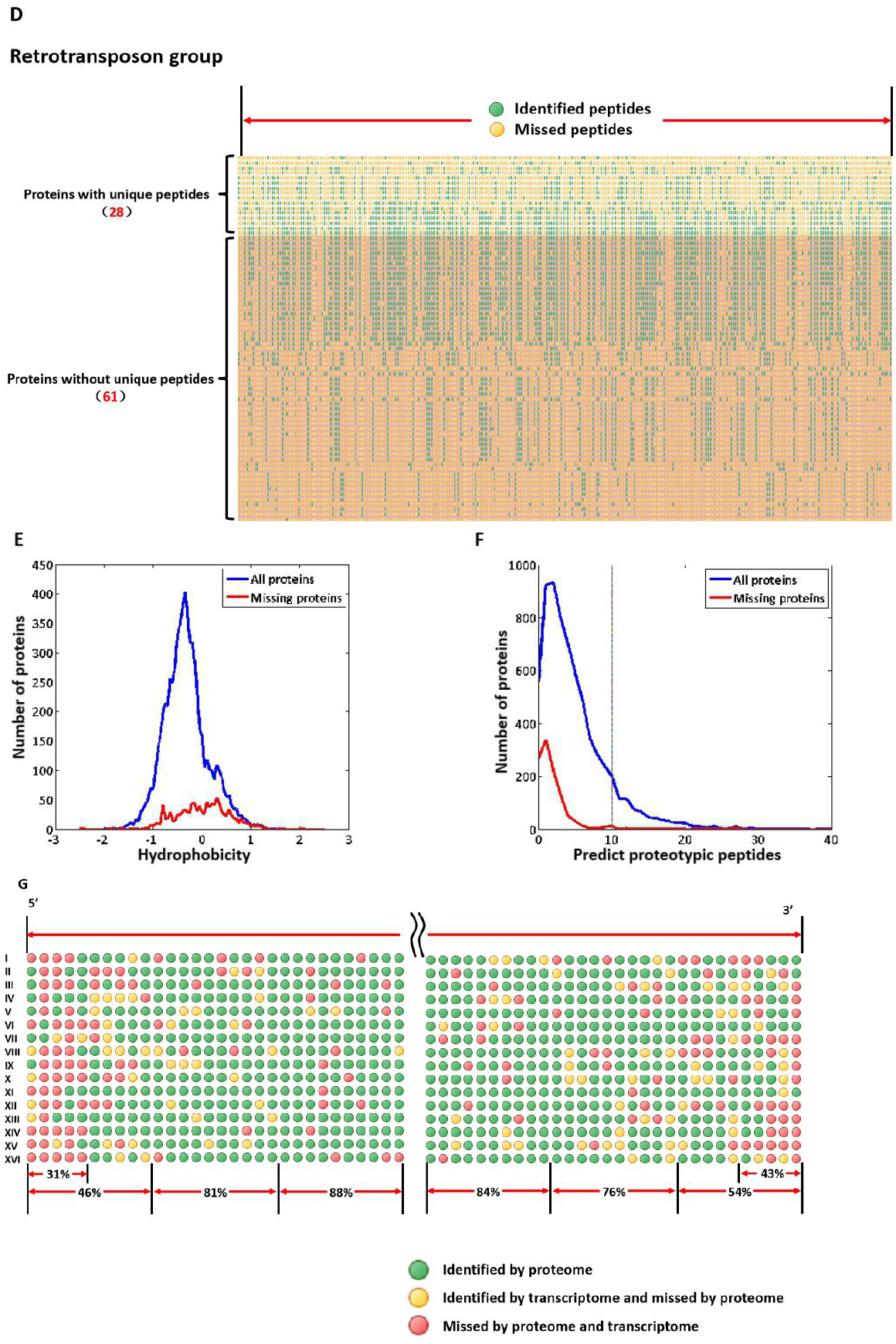
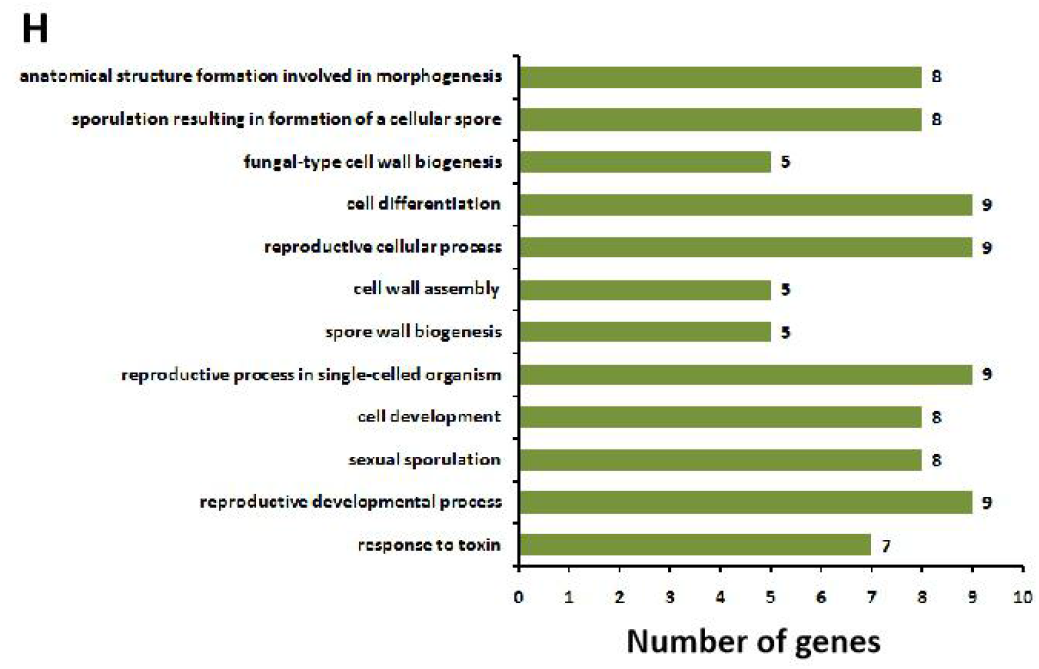
Missing proteins are heavily enriched for protein groups with high sequence homology. A, 149 proteins missed by our proteome dataset shared high-confidence peptides with the identified proteins. B, Classification of missing proteins with identified peptides. Protein with sequence coverage less than 10% would be signed as “no homology”. Three groups, retrotransposon, helicase, and ribosome, were found to be significantly enriched with conserved sequences. C, Visualization of the alignment of the sequenceable peptides for the protein group of helicase. 10 proteins were regarded as identified proteins for their unique peptides identification. 21 proteins were regarded as missing proteins for the absence of unique peptides. D, Visualization of the alignment of the sequenceable peptides for the protein group of retrotransposon. 28 proteins were regarded as identified proteins for their unique peptides identification. 61 proteins were regarded as missing proteins for the absence of unique peptides. E, Hydrophobicity distribution of missing proteins and all theoretical proteins. F, Distribution of the number of the predict proteotypic peptides among missing proteins and all theoretical proteins. Proteotypic peptides were predicted by PeptideSieve with threshold score larger than 80. G, Gene loci distribution of identified and missing proteins on chromosome. Green points represent the identified proteins in transcriptome and proteome. Yellow points represent the proteins identified by transcriptome but missed by proteome. Red points represent the proteins missed in both. Percentage represents the proportion of proteins identified by our proteome. H, Gene Ontology categories of biological processes of 44 missing proteins which have no significant characteristics on mRNA abundance, gene annotations, and protein physicochemical properties.

**Fig. S6.**
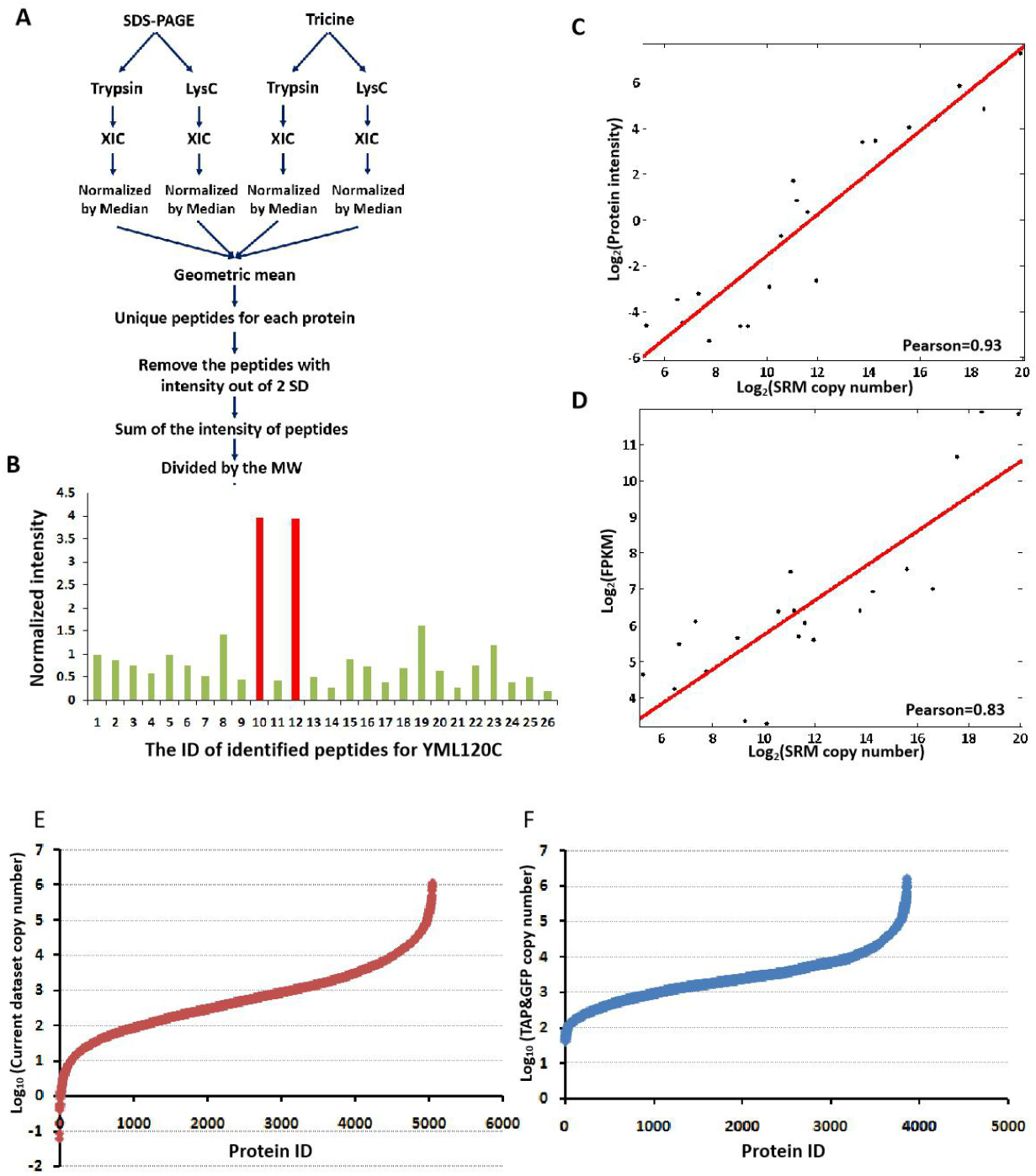
Dynamic range of our quantitative proteome based on label-free quantification analysis. A, Workflow for the normalization of label-free quantification of our proteome dataset. B, Normalized intensity of all identified peptides from YML120C. The red bar represents the peptide with abnormal intensity. C, Correlations between our quantified proteome and SRM datasets (Picotti *et al*., 2013). The *x*-axis represents the log_2_ protein copy number in SRM dataset, and the *y*-axis represents the log_2_ protein intensity in our quantitative proteome. D, Correlations between SRM dataset(Picotti *et al*., 2013) and transcriptome (Li *et al*., 2019). The *x*-axis represents the log_2_ protein copy number in SRM dataset, and the *y*-axis represents the log_2_ FPKM. E, Dynamic range of our quantified proteome. F, Dynamic range of TAP&GFP datasets(Ghaemmaghami *et al*., 2003; Huh *et al*., 2003).

**Fig S7.**
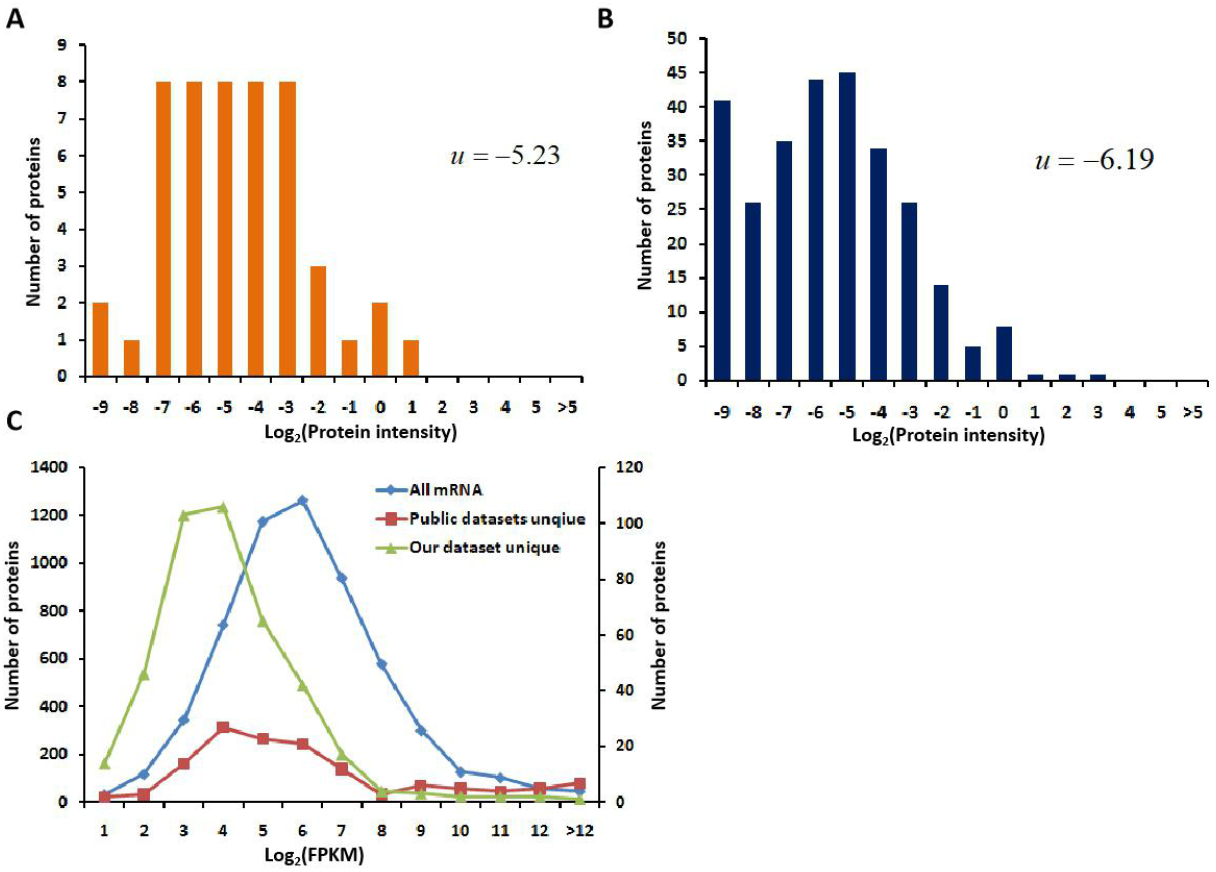
Intensity distribution of unique identified proteins in our proteome dataset. A, The intensity distribution of 241 unique proteins identified in our dataset vs RNA-seq dataset (Fig 3B). B, The intensity distribution of 609 unique proteins identified in our dataset vs four published datasets (Fig 2C). C, The distribution of unique proteins in our dataset (green line, right *y*-axis) (Fig 2C), uniquely in four published datasets (red line, right *y*-axis) (Fig 2C), and all proteins quantified by RNA-seq (blue line, left *y*-axis)(Fig 4B) based on mRNA abundance.

**Figure S8.**
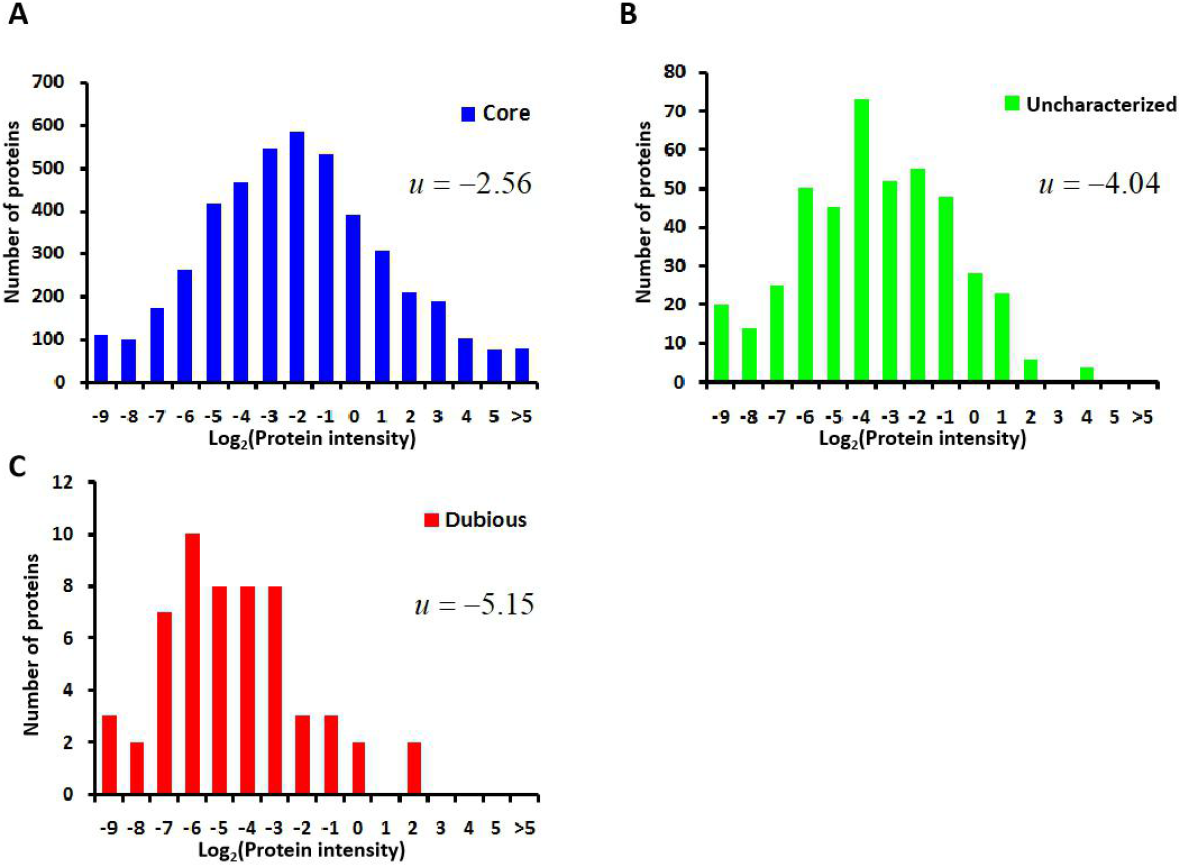
Intensity distribution of core proteins (A), uncharacterized proteins (B), and dubious proteins (C).

**Fig S9.**
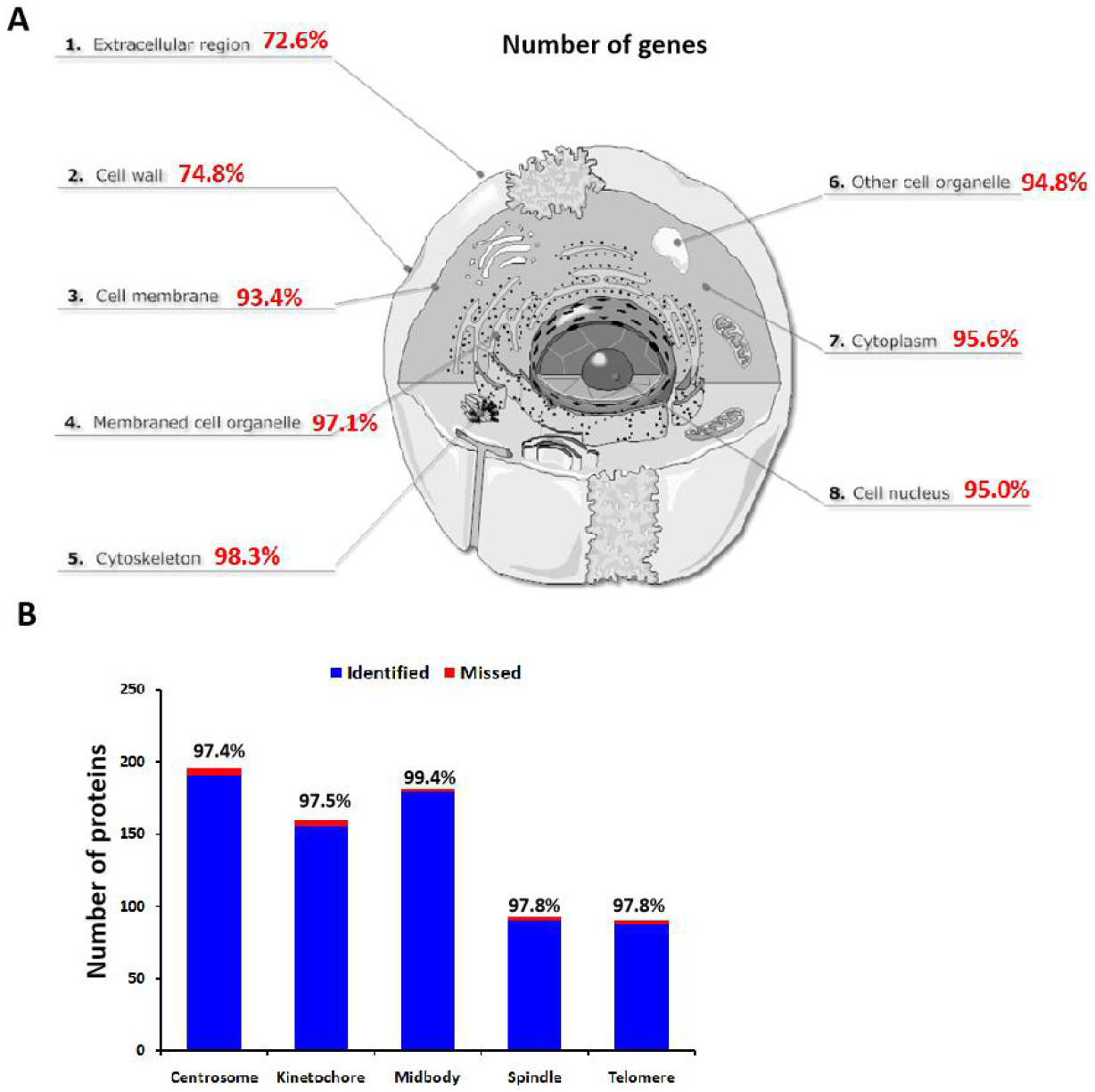
High coverage of all cellular components. A, Overview of proteome coverage in yeast cell. Percentage represents the proportion of identified proteins over the theoretical proteins in the given component of cell. B, Proteome coverage for five subgroups of mitosis proteins in yeast.

